# Varicella-zoster virus (VZV)-human interacting proteins enable the clustering of giant cell arteritis (GCA) samples: supports the association of VZV with a subset of GCA cases

**DOI:** 10.1101/2025.09.15.676295

**Authors:** Wei Ye, Xin yu Jiang, Feng Ben Shen

**Affiliations:** Department of Internal Medicine, Xiamen Susong Hospital, Xiamen, China; Department of Surgery, Women and Children’s Hospital, School of Medicine, Xiamen University, Xiamen, China; Beijing University of Technology, Beijing, China

**Keywords:** Varicella-Zoster Virus-Human Interaction Protein, Varicella-Zoster Virus, Giant Cell Arteritis, Spectral Clustering, Monte Carlo Simulation, Proteome, Transcriptome

## Abstract

**Objective:** To investigate the association between Varicella-Zoster Virus (VZV) and Giant Cell Arteritis (GCA).

**Methods:** One publicly available plasma proteomics dataset and three transcriptomics datasets were used. From the GCA samples, the corresponding values of VZV-human interacting proteins were extracted, and spectral clustering was performed on the GCA samples to calculate the silhouette coefficient. Subsequently, from the GCA samples, proteins or genes of the same quantity were randomly sampled 10,000 times; the identical spectral clustering method was applied to these randomly sampled sets, and the silhouette coefficient was calculated for each. Differences in spectral clustering performance and silhouette coefficients were compared between the VZV-human interacting proteins and the 10,000 randomly sampled protein/gene sets in the GCA samples. The same entire analytical process was repeated for the control group.

**Results:** In the GCA samples, the VZV-human interacting proteins exhibited a distinct clustering effect. Their silhouette coefficient was greater than that of most randomly sampled protein or gene sets, and this difference was statistically significant. No similar phenomenon was observed in the control group.

**Conclusion:** Varicella-Zoster Virus is associated with a subset of Giant Cell Arteritis cases.

## INTRODUCTION

Giant Cell Arteritis (GCA) is the most common systemic vasculitis in individuals aged over 50 years ^[1]^, with its incidence increasing steadily with age and being highest in Caucasians^[2]^. The typical symptoms of GCA include headache, jaw claudication, scalp tenderness, and features of Polymyalgia Rheumatica (PMR), such as pain in the shoulder and hip girdles ^[3]^. One of the most feared clinical complications of GCA is permanent visual loss, which may occur early in the disease course and is attributed to anterior ischemic optic neuropathy (AION), retinal artery occlusion, posterior ischemic optic neuropathy (PION), or cortical ischemia. Other complications of GCA include aortic aneurysm and dissection, cardiovascular events ^[4]^, and venous thromboembolic events ^[5]^.

Although the etiology of GCA remains to be fully elucidated, the dysregulated interaction between the vascular wall and the immune system appears to be crucial for its pathogenesis ^[6]^. Activated vascular dendritic cells recruit CD4+ T cells and macrophages to the vascular wall, which then enter through the blood vessels and migrate through the tissue spaces from the adventitia to the intima, leading to granuloma formation. Some studies suggest that the incidence of GCA has increased following the introduction of the live varicella-zoster vaccine ^[7]^.

Varicella-Zoster Virus (VZV) is a human herpesvirus. VZV can cause a range of vascular lesions, and pathological specimens of affected blood vessels exhibit features similar to those of GCA, such as multinucleated giant cells^[8]^. After detailed examination of temporal artery and aortic specimens from GCA patients with positive and negative biopsies, VZV antigens and DNA were detected in some studies^[9][10][11][12][13]^. However, multiple other studies failed to identify VZV in temporal artery specimens of GCA^[14][15][16][17][18][19][20][21][22]^.

A multicenter retrospective cohort study ^[23]^ evaluated the long-term risk of developing GCA after herpes zoster ophthalmicus (HZO). In this large real-world cohort, HZO was associated with an increased risk of developing GCA, although the absolute risk remained low.

In conclusion, the relationship between VZV and GCA is not yet fully clear. Through cluster analysis of publicly available proteomic and transcriptomic data, this study observed an important phenomenon regarding the association between VZV and some cases of GCA.

## METHODS

### (Selection of Target Human Proteins)

The VZV–human host protein interaction data analyzed in this study were derived from the high-confidence interactome results reported by Uetz et al. in the journal ⟪Science⟫ (detailed data see Supplementary Table S10 of this study,containing a total of 28 proteins) [Uetz et al., Science 311, 239–242, 2006; https://www.science.org/doi/10.1126/science.1116804].

A total of 28 core human target proteins were selected for subsequent cluster analysis, including ASF1B, ACADM, CDK3, CDC6, COPB2, CYB5D2, DCAF10, DNAJC21, EIF3M, H2AC6, H2AX, ORC2, ORC4, ORC5, PCNA, PPID, PRKACA, PRKACB, PRKAR2A, POLD2, PYGL, RAB7A, RRM1, RRM2, SAMHD1, STIM1, SUN2, WDFY2; these proteins are mainly involved in cell cycle regulation, DNA replication, and immune response pathways.

### (Data Source)

A total of 4 public datasets were used in this study, including plasma proteomics data and gene expression profile data, with detailed information as follows:

1. Plasma_GCA_2024 dataset^[24]^: A plasma proteomics dataset related to Giant Cell Arteritis (GCA). The dataset and corresponding research paper can be accessed via the following links: data link (https://github.com/cunni319/Plasma_GCA_2024), paper link [Cunningham KY, et al. Ann Rheum Dis 2024;0:1–11. doi:10.1136/ard-2024-225868].
2. GSE201753 dataset^[25]^: A gene expression profile dataset in the Gene Expression Omnibus (GEO) database. The dataset and corresponding research paper can be accessed via the following links: data link (https://www.ncbi.nlm.nih.gov/geo/query/acc.cgi?acc=GSE201753), paper link [Estupiñán-Moreno E, et al. Ann Rheum Dis 2022;81:1290–1300. doi:10.1136/annrheumdis-2022-222156].
3. Aortitis_2022 dataset^[26]^: An aortitis-related transcriptomics dataset. The dataset and corresponding research paper can be accessed via the following links: data link (https://github.com/jaeyunsung/Aortitis_2022), paper link [Arthritis Rheumatol. 2022 August; 74(8): 1376–1386. doi:10.1002/art.42138].
4. GSE174694 dataset^[27]^: A gene expression profile dataset in the GEO database. The dataset and corresponding research paper can be accessed via the following links: data link (https://www.ncbi.nlm.nih.gov/geo/query/acc.cgi?acc=GSE174694), paper link [Neurol Neuroimmunol Neuroinflamm 2021;8:e1078. doi:10.1212/NXI.0000000000001078].

### (Data Preprocessing)

Differentiated normalization strategies were adopted for different types of datasets (proteomics, transcriptomics) to eliminate data heterogeneity, with specific steps as follows:

Proteomics datasets (Plasma_GCA_2024) : First, quantile normalization was used to standardize the data between samples to ensure consistent protein expression distribution across different samples; subsequently, gene (protein)-wise Z-score normalization was performed on the normalized protein expression data—for the expression values of each protein across all samples, the mean and standard deviation were calculated, and then the expression values were converted to Z-scores using the formula (Z = (x - μ)/σ, where x is the original expression value, μ is the mean, and σ is the standard deviation), so that the expression of each protein follows a standard normal distribution among samples.

Transcriptomics datasets (GSE201753, Aortitis_2022, GSE174694) : First, the DESeq2 package was used to perform variance-stabilizing transformation (VST) on the original count data to reduce data heteroscedasticity and retain biological differences between samples; subsequently, gene-wise Z-score normalization was performed on the VST-transformed data, with the same normalization formula as that for proteomics data, to ensure the comparability of gene expression data in subsequent analyses.

### (Spectral Clustering Analysis)

Spectral clustering analysis was performed on the preprocessed target protein/gene expression data to explore the grouping patterns among samples, with the specific process as follows:

1. Distance matrix calculation: The expression matrix of target proteins/genes was transposed (to ensure rows represent samples and columns represent molecules), and then the Euclidean distance matrix between samples was calculated based on the transposed matrix.
2. Affinity matrix construction: The 50th quantile of the distance matrix was used as the sigma (σ) parameter, and the affinity matrix between samples was constructed via the Gaussian kernel function (affinity = exp(-dist^2^/(2σ^2^))); the diagonal elements of the affinity matrix were set to 0 (to exclude the association of a sample with itself); if the row sum of a certain row in the affinity matrix was 0, all elements in this row were set to 1e-10 to avoid errors in subsequent calculations.
3. Laplacian matrix calculation: First, the row sum of each row in the affinity matrix was calculated to construct a diagonal matrix D (diagonal elements are row sums, others are 0), and the inverse square root transformation was performed on D (D_inv_sqrt = diag(1/sqrt(row_sums))); subsequently, the normalized Laplacian matrix was calculated via the formula (Laplacian = I - D_inv_sqrt × affinity × D_inv_sqrt, where I is the identity matrix).
4. Eigenvalue and eigenvector extraction: Eigenvalue decomposition was performed on the Laplacian matrix, eigenvalues were sorted in ascending order, and the first k eigenvectors (k=2 in this study) corresponding to the sorted eigenvalues were extracted to form matrix U; matrix U was normalized (each row element divided by the square root of the sum of squares of elements in that row) to obtain matrix Y.
5. K-means clustering: Using matrix Y as input, the k-means clustering algorithm (number of clusters k=2, number of random initial cluster center selections nstart=25) was used to cluster samples, and the final sample clustering results were obtained.

### (Permutation Test Validation)

To verify the statistical significance of the clustering results of target proteins/genes, a permutation test was used to compare the clustering effect of target molecules with that of random molecules, and the fairness of the test was strictly ensured, with specific steps as follows:

1. Fairness design: In the permutation test, the number of randomly selected molecules (proteins/genes) was completely consistent with that of target molecules; in terms of data source, random molecules were all derived from the full set of molecules in the corresponding analysis group (same as the data source of target molecules); in terms of clustering and evaluation methods, the spectral clustering parameters (e.g., sigma calculation method, k value, nstart times) and silhouette coefficient calculation method for random molecules were completely consistent with those for target molecules, to eliminate the interference of methodological differences on results.
2. Permutation process: For each analysis group, random molecule extraction and clustering analysis were repeated 10000 times; after extracting random molecules each time, the same spectral clustering process as that for target molecules was used to obtain clustering results, and the silhouette coefficient of the corresponding cluster was calculated (implemented via the ‘cluster’ package, with the formula sil_width = (b - a)/max(a, b), where a is the average distance between the sample and other samples in the same cluster, and b is the average distance between the sample and samples in different clusters).
3. Significance calculation: The number of times the silhouette coefficient of random molecule clustering was greater than or equal to that of target molecule clustering in 10000 permutations was counted; the ratio of this number to the total number of permutations (10000) was used as the P-value, and a P-value < 0.05 indicated that the clustering effect of target molecules was statistically significant.

### (Specification of Sigma Parameter)

The sigma (σ) parameter used for affinity matrix construction in spectral clustering in this study was calculated based on the quantile of the inter-sample distance matrix; verification showed that when the quantile value varied within the range of 0–1, the change of the sigma parameter had little impact on the final clustering results (e.g., sample clustering grouping, silhouette coefficient), and the clustering effects under different quantiles were consistent.

## RESULTS

### Introduction to the Plasma_GCA_2024 Dataset and Spectral Clustering Analysis

The study cohort included 30 patients with biopsy- or imaging-confirmed giant cell arteritis (GCA), along with 30 age-matched, sex-matched, and race-matched non-disease controls. Plasma samples were collected from all participants. Active disease was defined as having a Physician Global Assessment (PGA) score ≥ 2. Clinical remission was defined as having a PGA score of 0, an ESR < 30 mm/hour, and a CRP level < 10 mg/L. Plasma proteome profiling was performed using the SomaScan Assay version 4 from SomaLogic.

**Figure 1.**
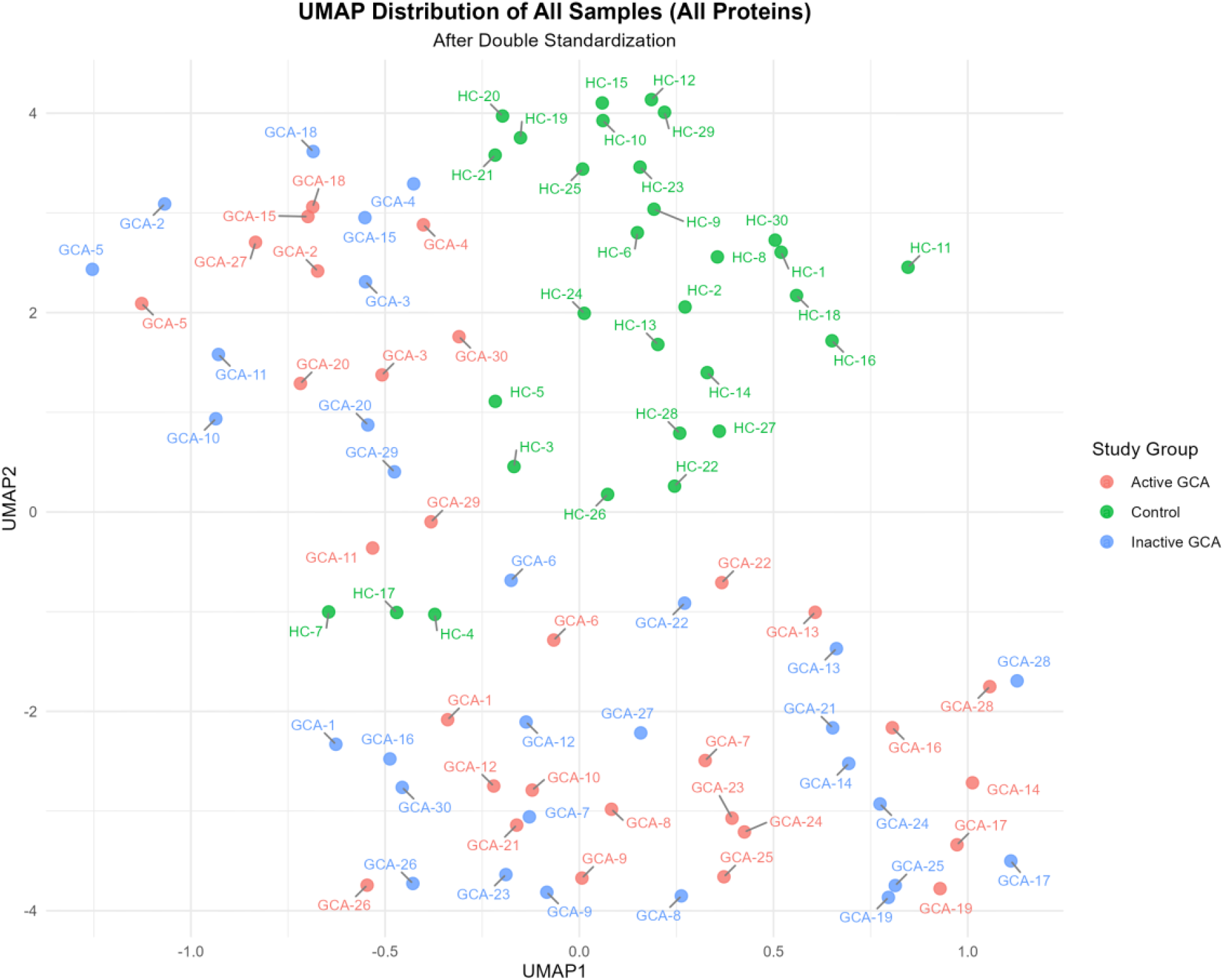
UMAP Distribution of All Samples Based on All Genes (After Double Standardization) Shows the UMAP dimensionality reduction distribution results of all samples (including Active Giant Cell Arteritis (Active GCA), Inactive Giant Cell Arteritis (Inactive GCA), healthy controls (Control), and other samples) based on all genes after double standardization.

**Figure 2.**
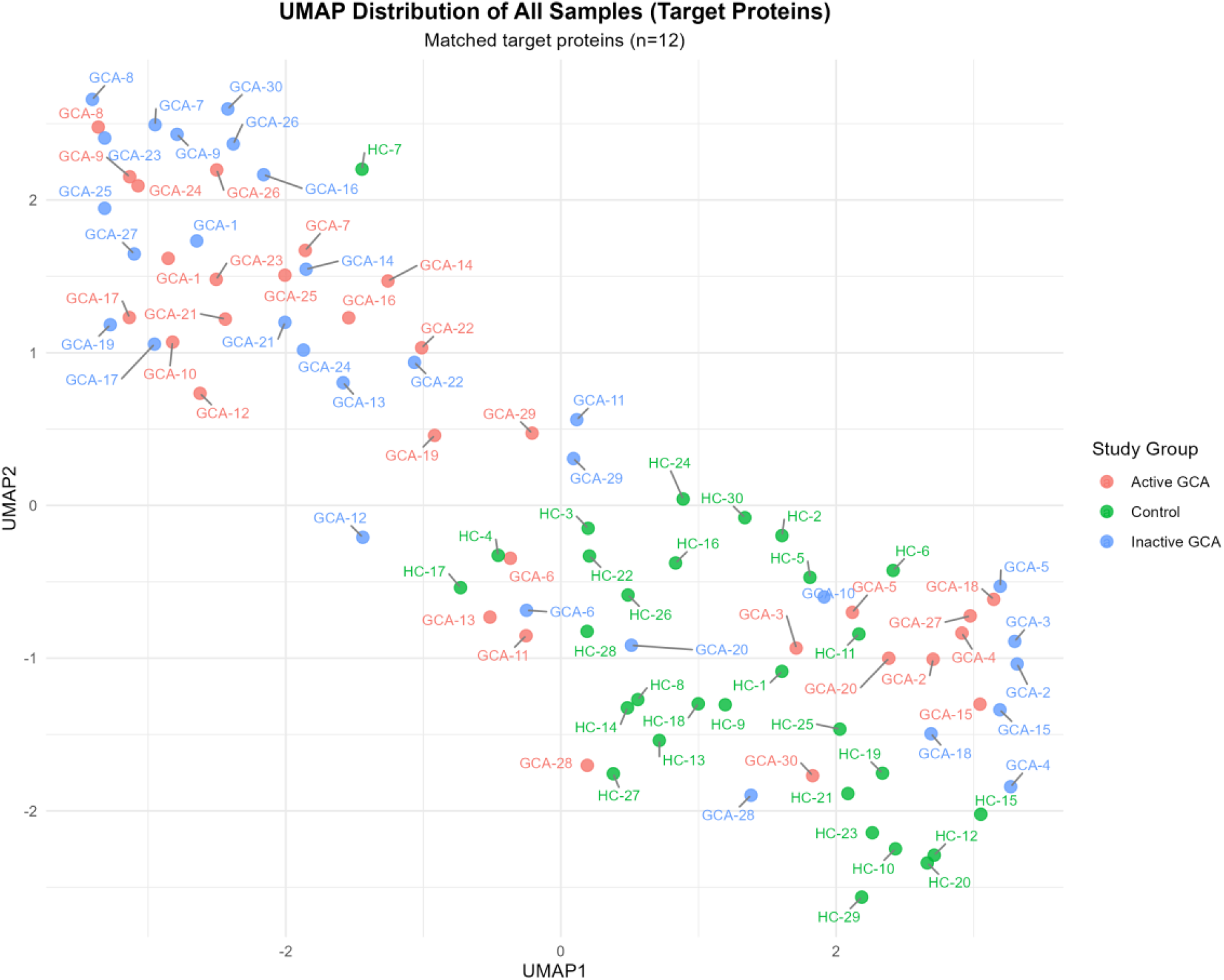
UMAP Distribution of All Samples Based on Target Proteins Shows the UMAP dimensionality reduction distribution results of all samples (including Active Giant Cell Arteritis (Active GCA), Inactive Giant Cell Arteritis (Inactive GCA), healthy controls (Control), and other samples), with 12 matched target proteins.

**Table 3.**
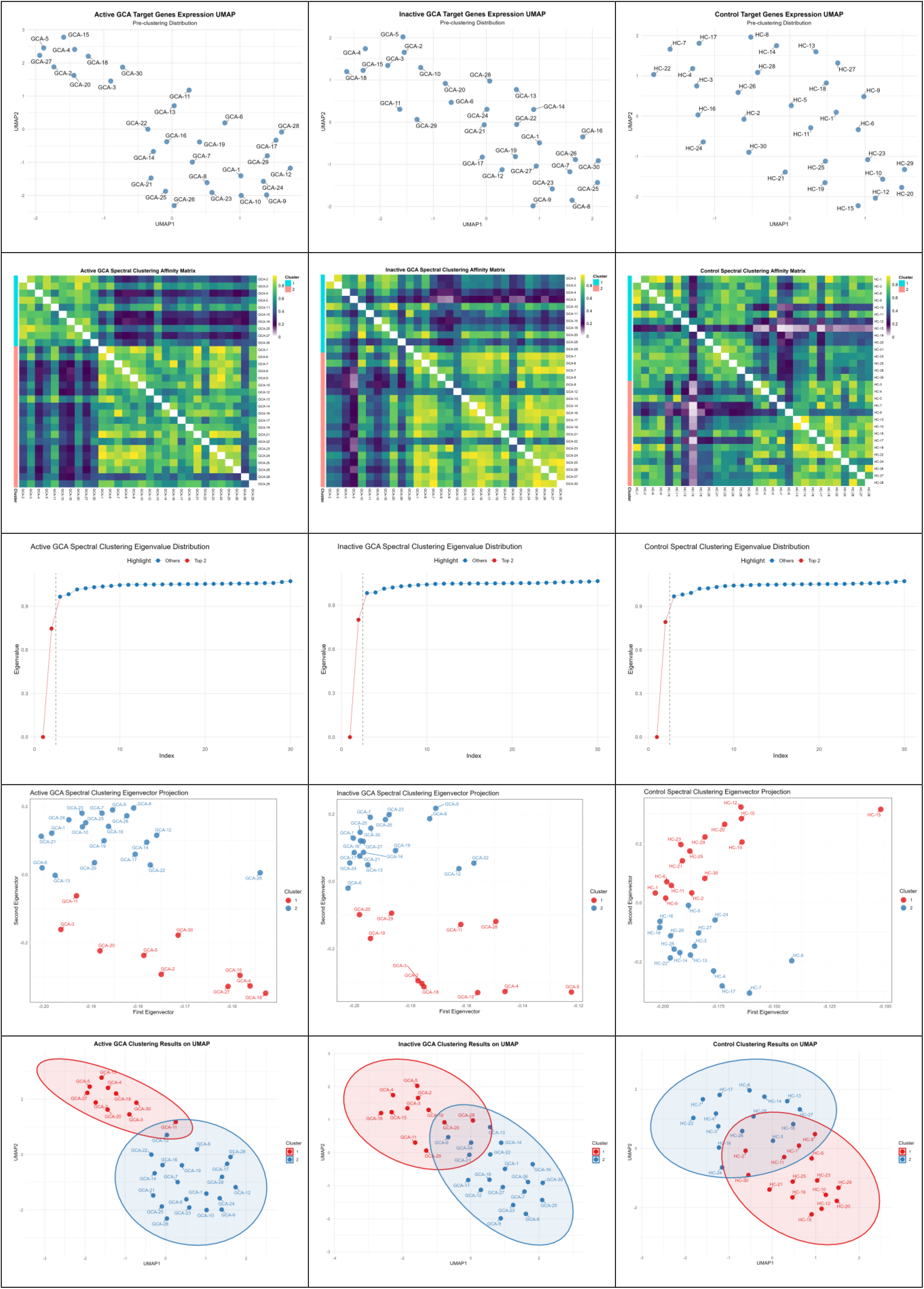
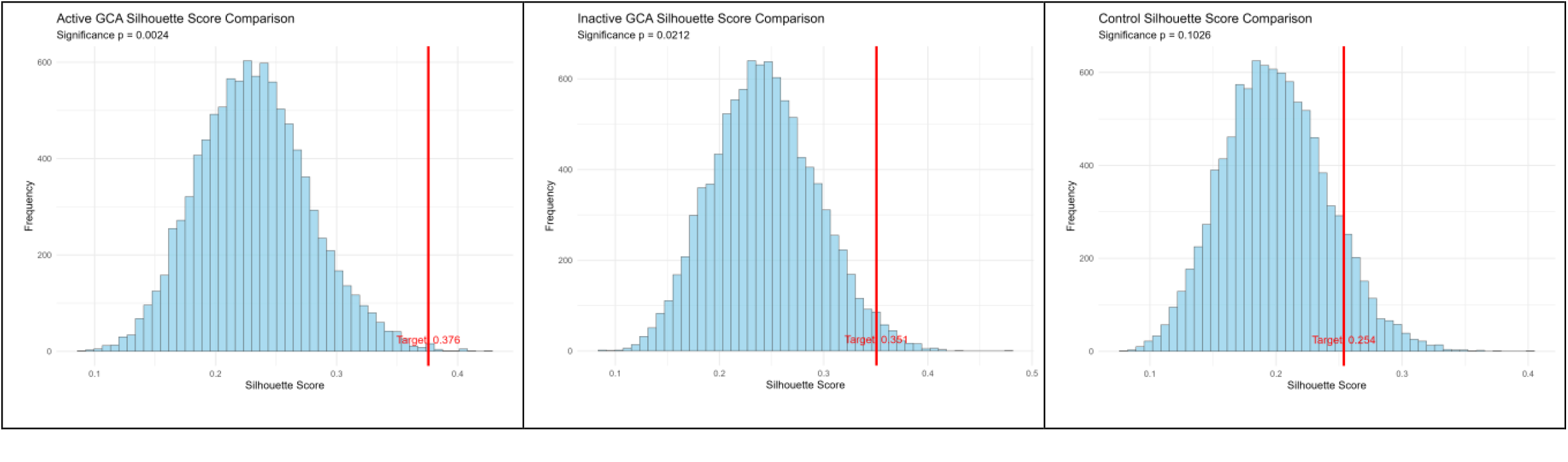
Spectral Clustering Analysis and Related Visualization Results for Different Study Groups.

Includes multiple key analysis results for the Active Giant Cell Arteritis (Active GCA), Inactive Giant Cell Arteritis (Inactive GCA), and healthy control (Control) groups: spectral clustering affinity matrix, spectral clustering eigenvalue distribution (highlighting the top 2 eigenvalues), spectral clustering eigenvector projection, clustering results in UMAP space, and the comparison of silhouette scores based on target genes in the Active GCA group (significance p=0.0024, silhouette score for target genes=0.376).

From the normalized database (which has the same data source as the target protein data), we randomly select a number of proteins equal to the number of valid target proteins to form a protein set, and then perform spectral clustering using the exact same method. This random selection process is repeated 10,000 times, generating 10,000 random protein sets. The spectral clustering results of these 10,000 random protein sets form a null distribution. The main purpose is to determine whether the target protein set is specific by comparing it with the random protein sets.

### Introduction to the GSE201754 Dataset and Spectral Clustering Analysis

The transcriptome analysis samples in this study were CD14+ monocytes, derived from 82 patients with Giant Cell Arteritis (GCA) and 31 healthy controls. The 82 GCA patients were cross-sectionally classified into three subgroups: (1) Active disease (ACT); (2) Remission with glucocorticoid (GC) treatment(RT),; (3) Remission without GC treatment(RNT).RNA Extraction: Total RNA was extracted from isolated and purified CD14+ monocytes to ensure the purity and integrity of RNA, laying the foundation for subsequent transcriptome analysis. RNA Sequencing: Transcriptome sequencing (RNA Sequencing, RNA-seq) was performed on the extracted total RNA, and genome-wide gene expression data of CD14+ monocytes were obtained using high-throughput sequencing technology.

**Figure 4.**
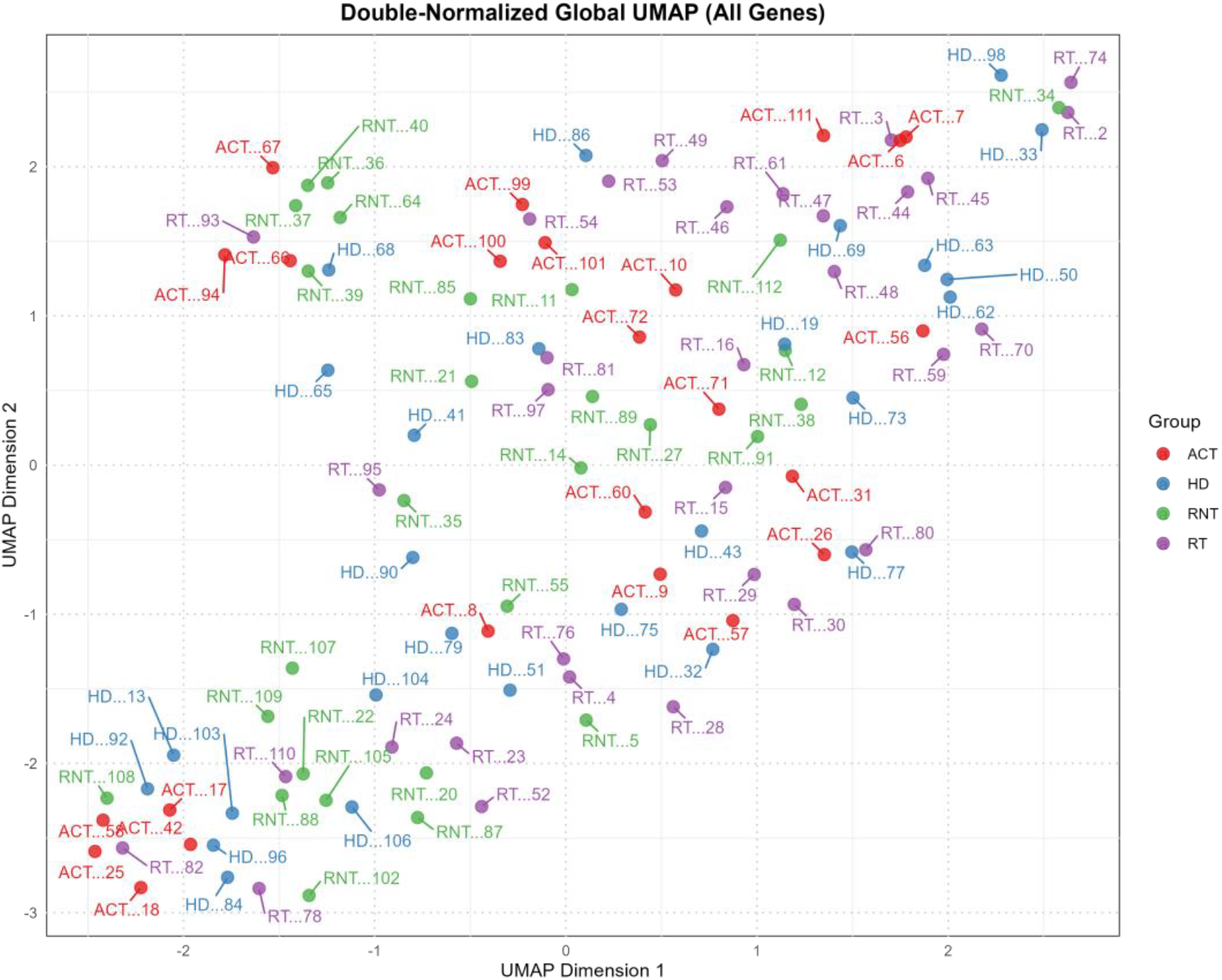
This figure is the double-normalized global UMAP (Uniform Manifold Approximation and Projection) dimensionality reduction visualization result based on all genes. It displays the distribution of samples from four experimental groups (ACT, RT, RNT, HD) in a two-dimensional space (with UMAP Dimension 1 as the horizontal axis and UMAP Dimension 2 as the vertical axis). Each sample is labeled with “group name + number” (e.g., HD…98, RT…74). Through the aggregation degree of samples in the space, the similarity of samples within different experimental groups and the difference degree of samples between groups can be intuitively observed, providing a global sample distribution reference for subsequent clustering analysis.

**Figure 5.**
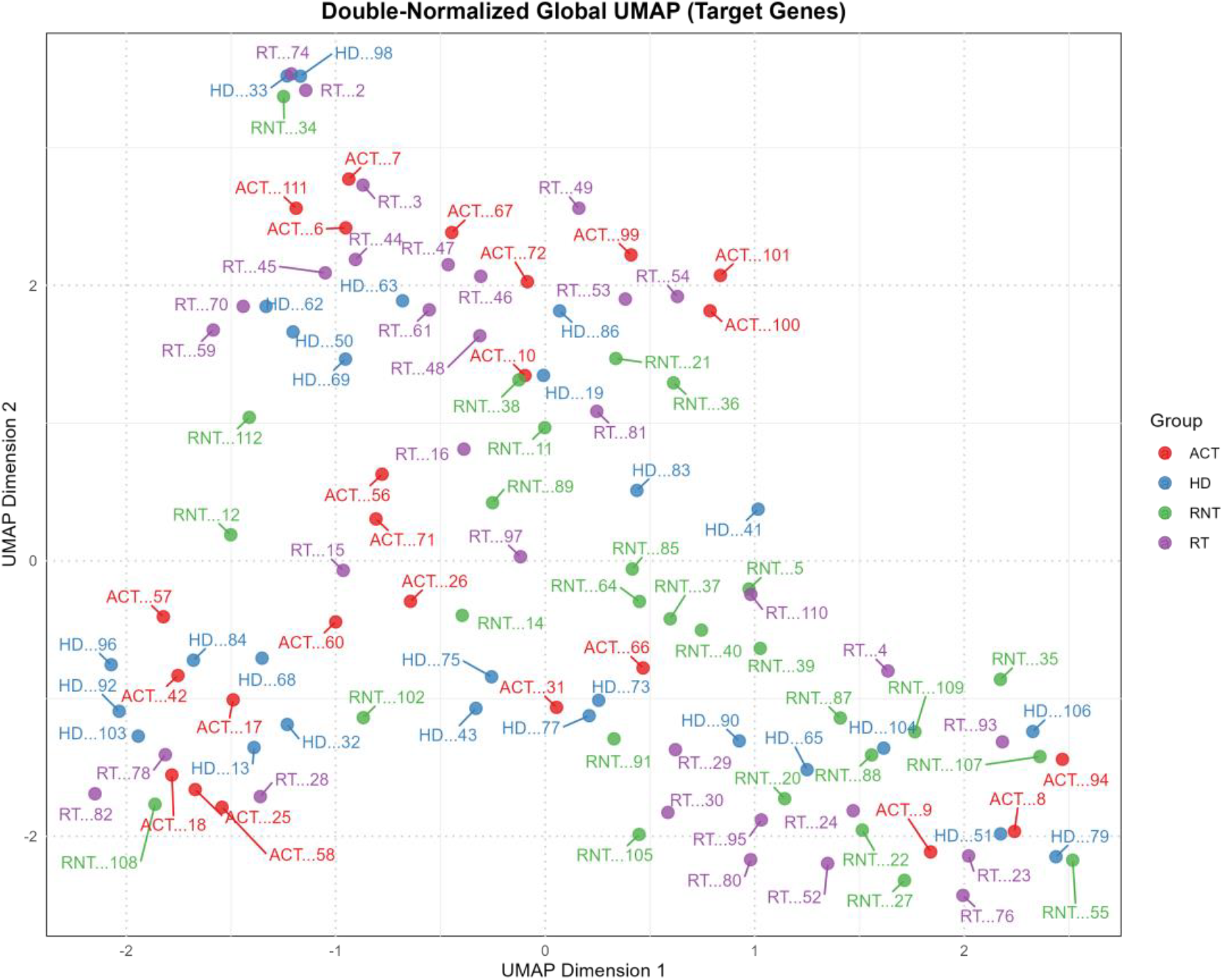
Double-Normalized Global UMAP Plot (Based on Target Genes) This plot is constructed after double normalization of expression data from target genes, with the horizontal axis representing UMAP Dimension 1 (range: -2∼2) and the vertical axis representing UMAP Dimension 2 (range:-2∼2); each point in the plot represents one sample, labeled with group information (HD, RT, ACT, RNT) and sample numbers; compared with the “global UMAP plot based on all genes”, this plot can focus more on the sample expression differences related to target genes, and is used to display the clustering distribution differences of samples from different groups under the expression pattern of target genes, providing a sample clustering.

**Table 6.**
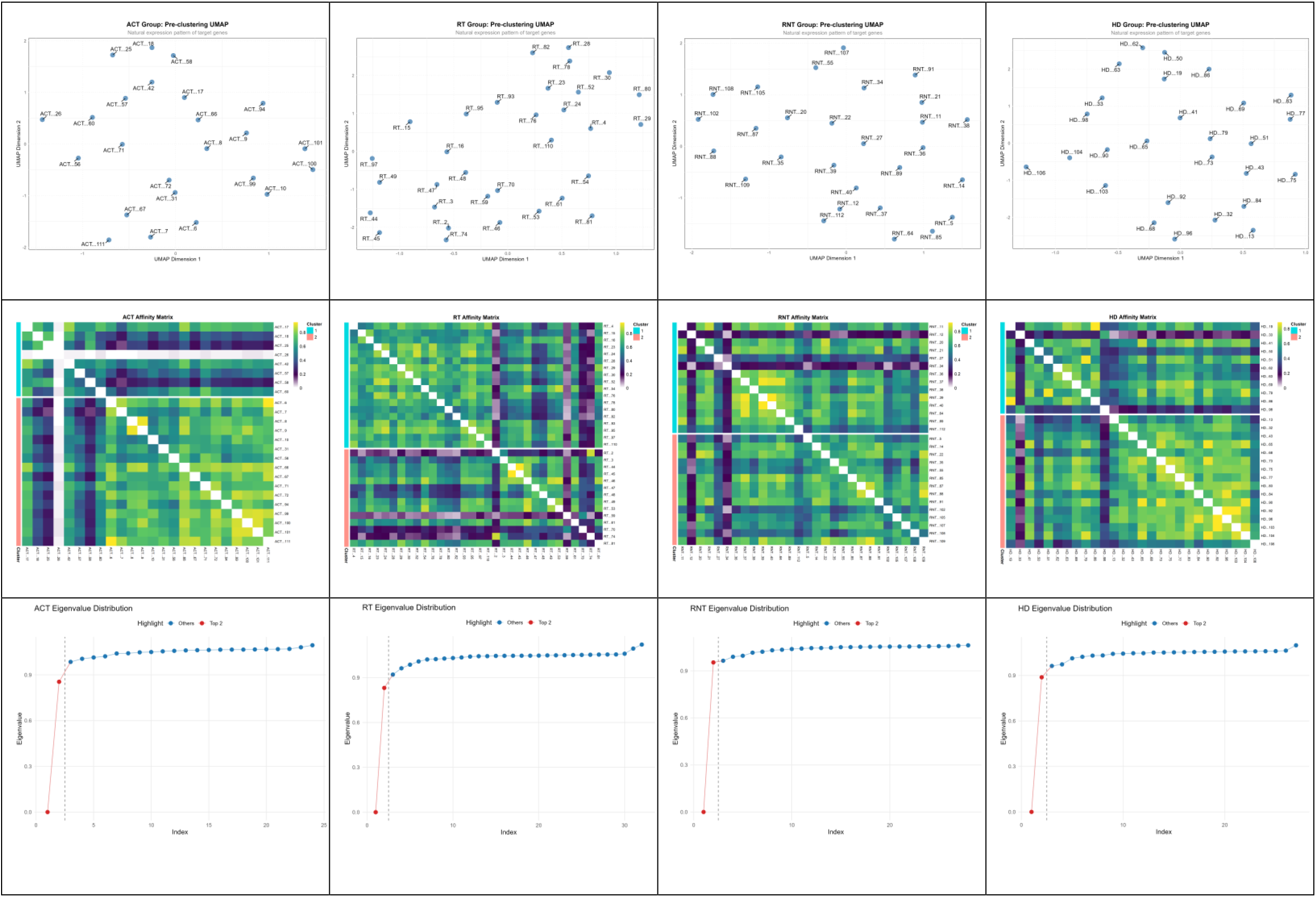
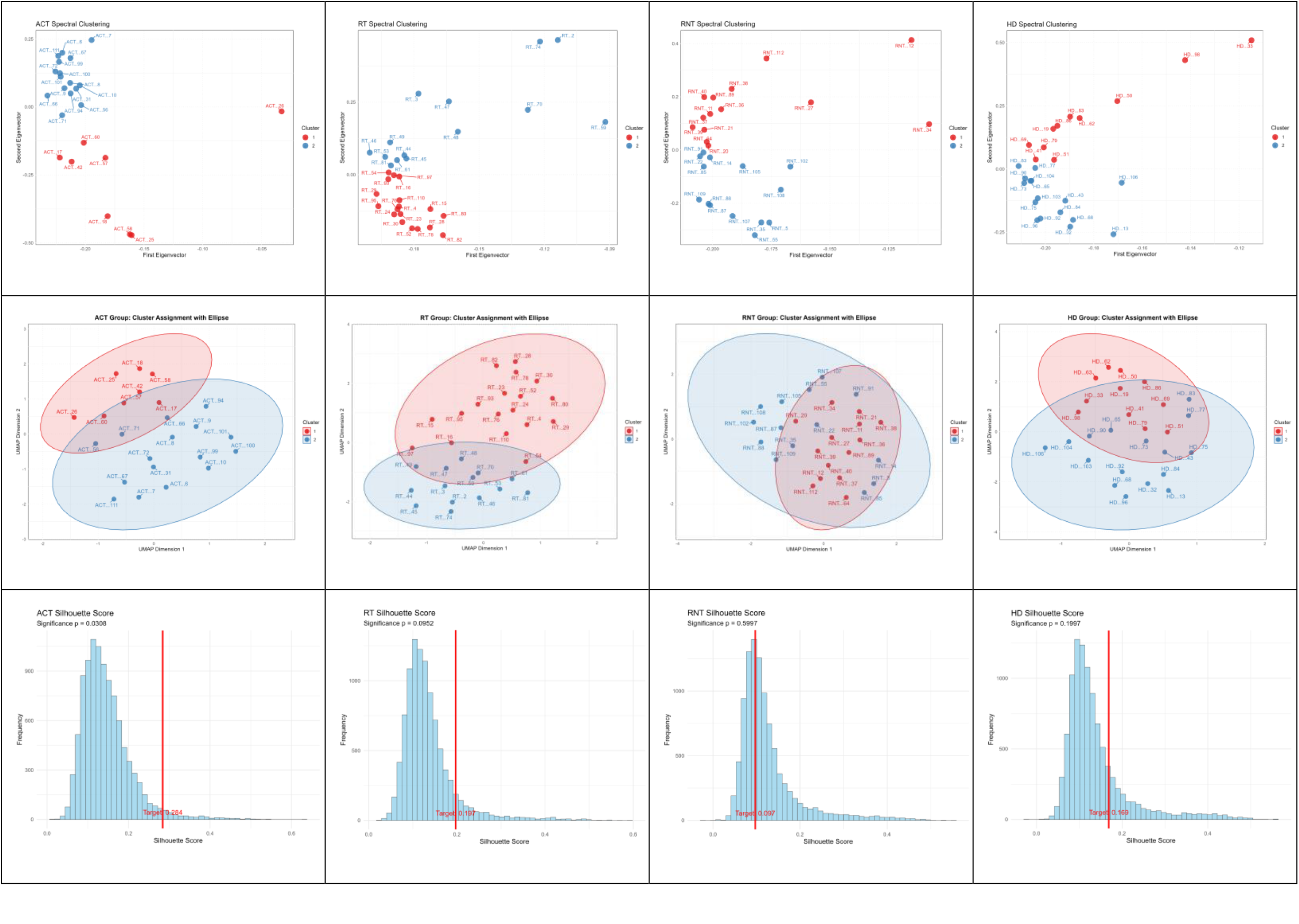
Multi-Dimensional Clustering Analysis Results of Samples in Different Groups (ACT, RT, RNT, HD)

This table integrates 6 types of key analysis plots for samples in 4 groups, including pre-clustering UMAP, affinity matrix, eigenvalue distribution, spectral clustering results, cluster assignment UMAP, and silhouette score. It comprehensively evaluates the clustering effect of samples in each group from qualitative (distribution, similarity) and quantitative (eigenvalue, silhouette score) perspectives.

### Introduction to the Aortitis_2022 Dataset and Spectral Clustering Analysis

The study samples were divided into an inflammatory aortic aneurysm group (n=25) and a non-inflammatory aortic aneurysm group (n=25), with the two groups matched in age and sex; the inflammatory aortic aneurysm group was further subdivided into patients with giant cell arteritis (GCA)/polymyalgia rheumatica (PMR) (n=8, with documented diagnosis or clinical features compatible with GCA and/or PMR) and patients with clinically isolated aortitis (CIA) (n=17, inflammatory aortitis without diagnosis of or clinical features compatible with GCA or PMR). The inclusion criterion for inflammatory aortic aneurysm samples (including clinically isolated aortitis) was the presence of ‘active giant cell aortitis’ in the pathological description of the resected ascending aortic tissue; patients with signs of localized or systemic infection were excluded by manual review of medical records; non-inflammatory aortic aneurysm samples were from patients who underwent surgical resection during the same period and were age- and sex-matched with the inflammatory group. Formalin-fixed paraffin-embedded (FFPE) blocks containing ascending aortic aneurysm tissues were cut into 10 μm-thick sections. Sequencing: The final libraries were quantified by Agilent TapeStation D1000 and Qubit dsDNA BR Assay Kit, and 101-bp paired-end reads were sequenced on an Illumina HiSeq4000 platform; samples of inflammatory and non-inflammatory aortic aneurysms were not sequenced separately, thus eliminating the need for batch correction protocols.

**Figure 7.**
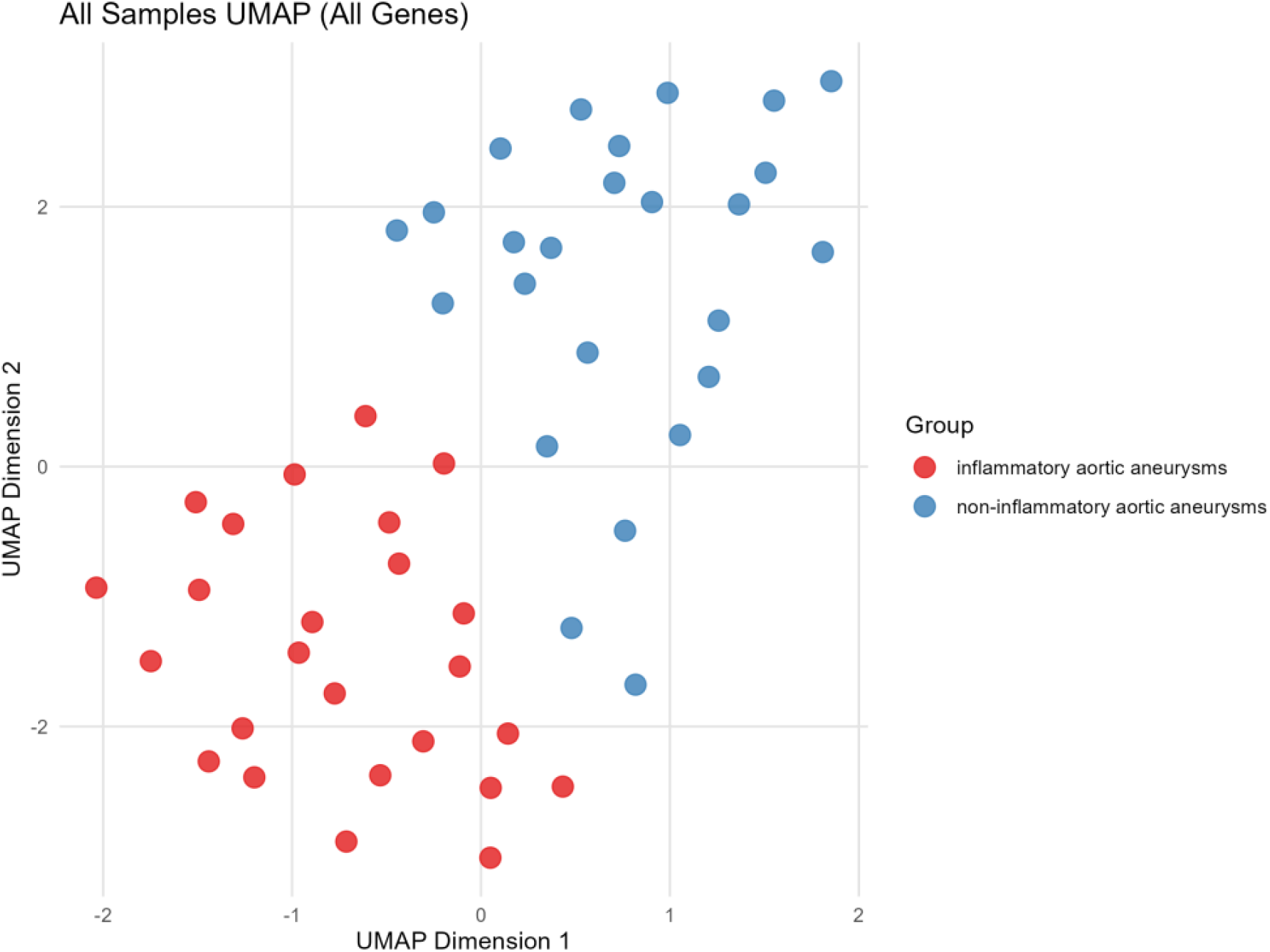
UMAP dimensionality reduction analysis plot of all samples based on all genes This figure uses the Uniform Manifold Approximation and Projection (UMAP) dimensionality reduction algorithm to present the distribution characteristics of all samples at the whole-gene expression level, and can clearly distinguish two types of samples: inflammatory aortic aneurysms (IAA) and non-inflammatory aortic aneurysms (NIAA).

**Figure 8.**
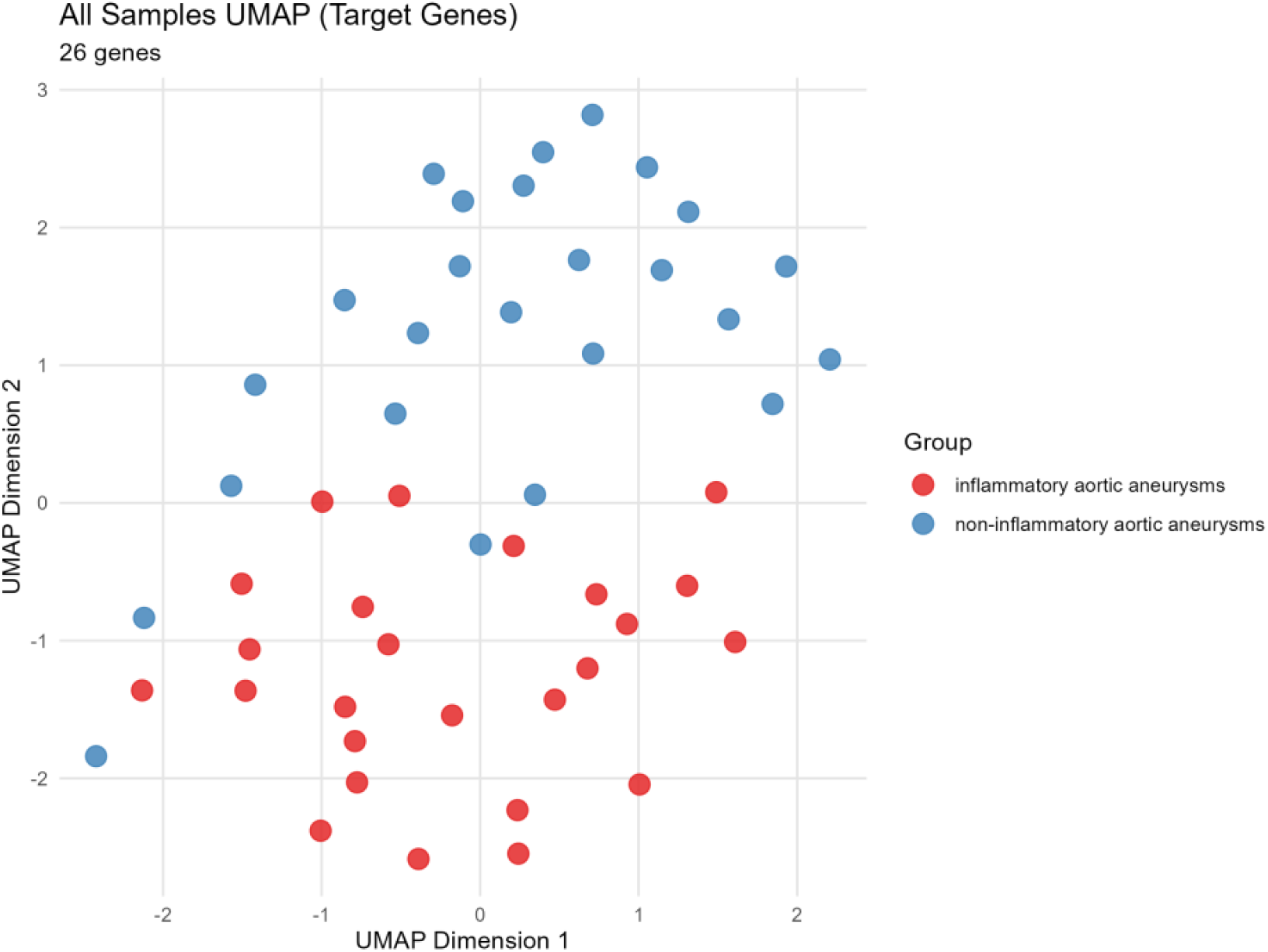
UMAP dimensionality reduction analysis plot of all samples based on 26 target genes Based on the expression data of 26 screened target genes, this figure uses the UMAP dimensionality reduction method to show the clustering distribution of all samples, and intuitively reflects the differences in target gene expression patterns between inflammatory aortic aneurysm (IAA) and non-inflammatory aortic aneurysm (NIAA) samples.

**Table 9.**
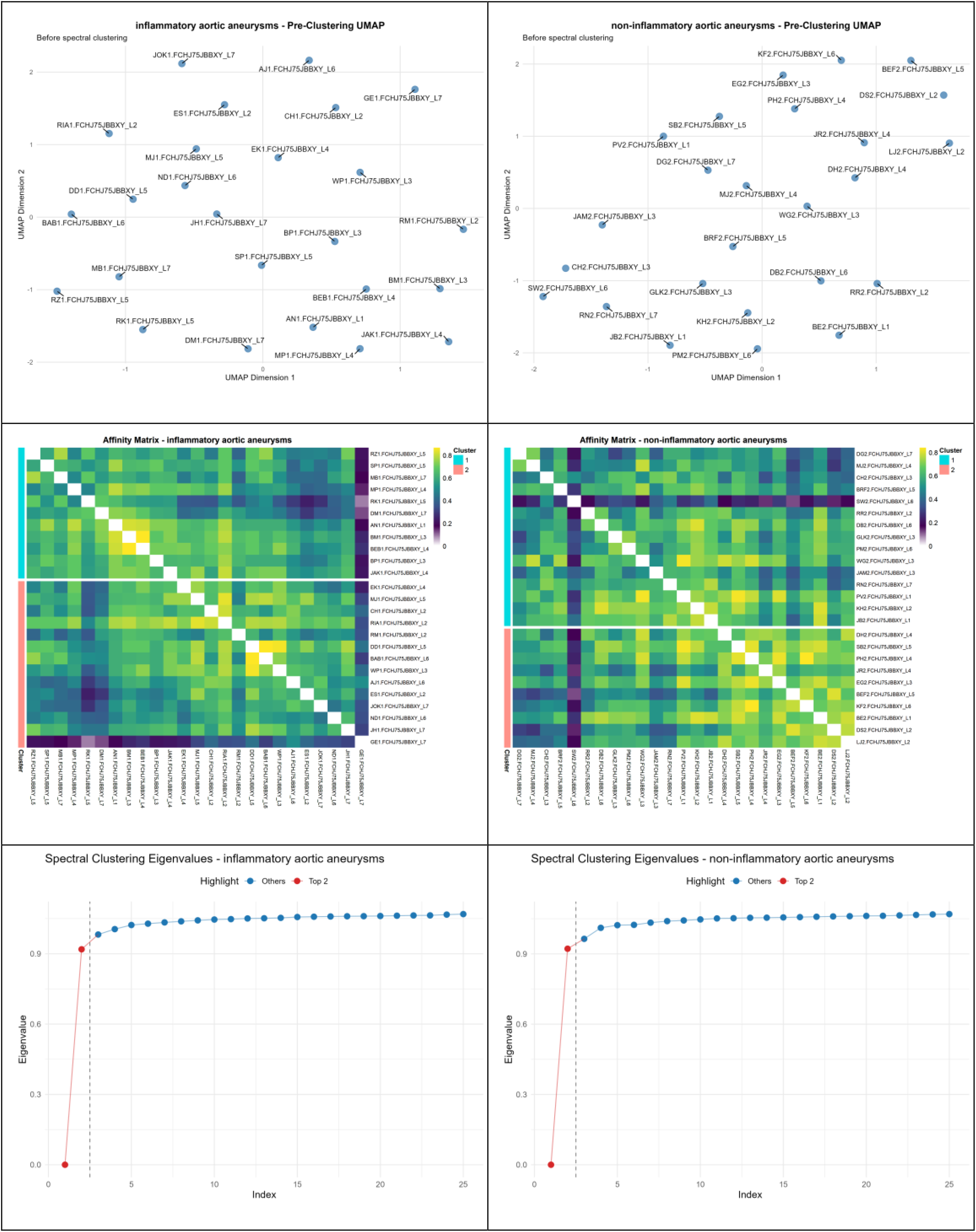
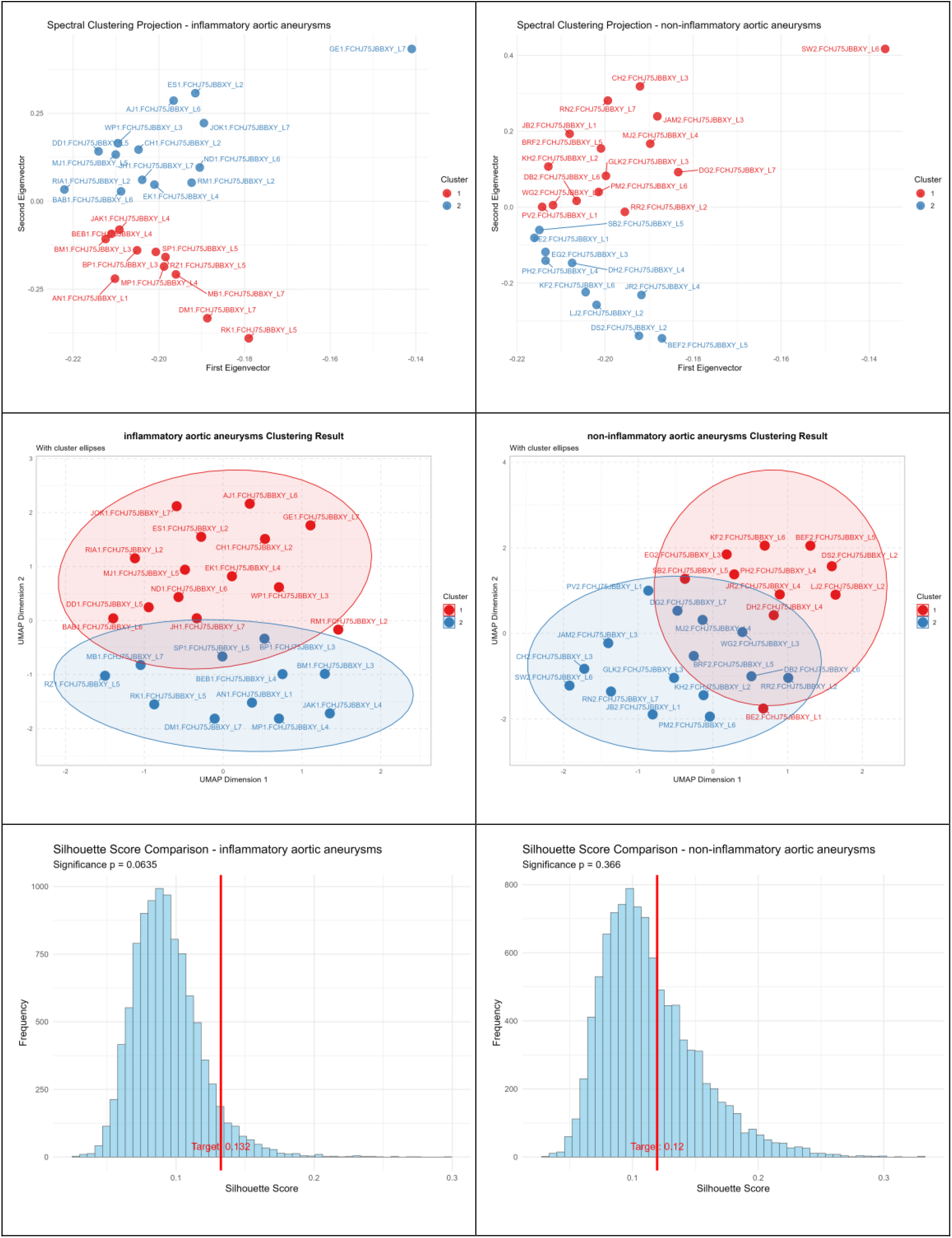
Summary table of multi-dimensional analysis results of inflammatory and non-inflammatory aortic aneurysm samples.

This table integrates multiple key analysis results of two types of aortic aneurysm samples, including multiple subplots: sample UMAP visualization based on target gene sets (presenting the characteristics of sample clustering distribution), affinity matrix of non-inflammatory aortic aneurysm samples (reflecting the degree of similarity between samples), spectral clustering projection of non-inflammatory aortic aneurysm samples (presenting clustering results based on the first eigenvector), and silhouette score comparison of inflammatory aortic aneurysm samples (silhouette score=0.132, significance P-value=0.0635, used to evaluate clustering effect).

*** Special Note: Although statistically non-significant (silhouette score=0.132, P=0.0635), the association between VZV and GCA in inflammatory aortic aneurysm specimens demonstrated borderline significance with consistent effect direction, suggesting potential biological relevance.

### Introduction to the GSE174694 Dataset and Spectral Clustering Analysis

Study Group (Giant Cell Arteritis, GCA): This study used 40 formalin-fixed, paraffin-embedded (FFPE) temporal artery biopsy sections from biopsy-positive giant cell arteritis (GCA) participants over 50 years of age. GCA cases were further divided into two subgroups: 20 contained varicella zoster virus (VZV) antigen (GCA/VZV-positive); the other 20 did not contain VZV antigen (GCA/VZV-negative). Control Group (Normal): This study used 31 normal FFPE temporal artery sections as controls, removed postmortem from participants over 50 years of age with no history of zoster, diabetes, cancer, substance abuse, or immunosuppression. Control group was also further divided into two subgroups: 11 contained VZV antigen (Control/VZV-positive); 20 did not contain VZV antigen (Control/VZV-negative). All control samples showed no morphological signs of inflammation. All GCA sections displayed classic biopsy-positive GCA features: transmural inflammation, medial damage, and multinucleated giant/epithelioid cells. Targeted RNA Sequencing (Targeted RNA Sequencing): FFPE temporal artery sections were analyzed using the TempO-Seq platform (BioSpyder Technologies, Carlsbad, CA) for targeted RNA sequencing of the whole human transcriptome. For samples with detected VZV antigen, tissues were scraped from the regions corresponding to VZV antigen-positive areas on unstained adjacent sections (10 mm×10 mm pooled regions of interest per participant) and placed into PCR tubes containing 1× lysis buffer; for samples without detected VZV antigen, tissues from equivalent areas of arteries on unstained sections were scraped.

**Figure 10.**
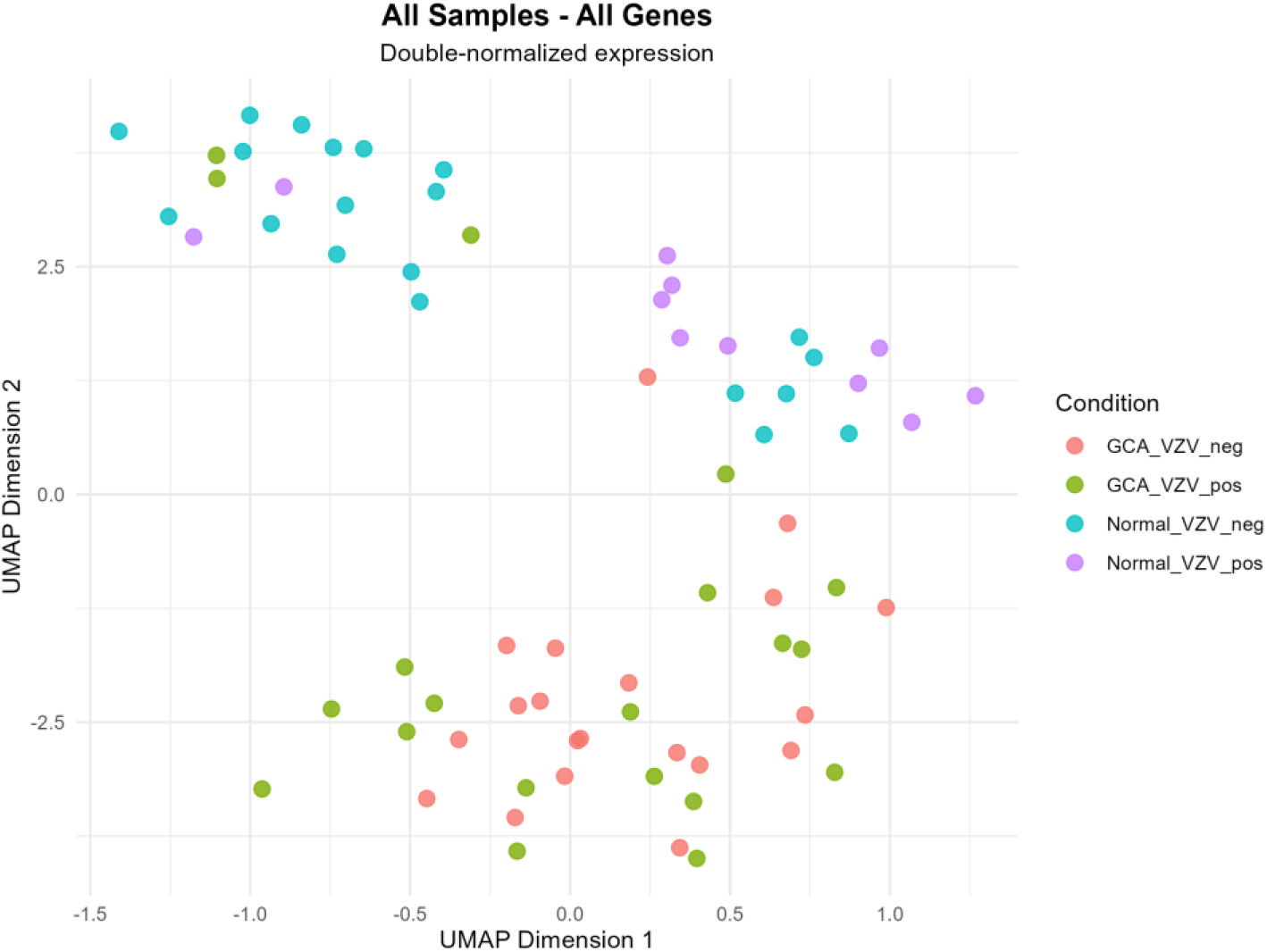
UMAP visualization of all genes in all samples (based on double-normalized expression). The horizontal axis is UMAP Dimension 1, and the vertical axis is UMAP Dimension 2; samples are divided into four groups according to experimental conditions, namely GCA_VZV negative group (GCA_VZV_neg), GCA_VZV positive group (GCA_VZV_pos), normal VZV negative group (Normal_VZV_neg) and normal VZV positive group (Normal_VZV_pos), and samples in different groups are distinguished by differential markers. ***Special Note: For this dataset, due to the large distance between samples, the possibility of batch effects must be considered. A different normalization strategy from the above three datasets has been adopted: the ‘blind’ parameter in the code vsd <-varianceStabilizingTransformation(dds, blind = TRUE) is adjusted from FALSE to TRUE; meanwhile, the operation scope of z-score is changed from all samples to the two groups of samples in paired clustering. The main purpose of these adjustments is to partially reduce the impact of batch effects.

**Figure 11.**
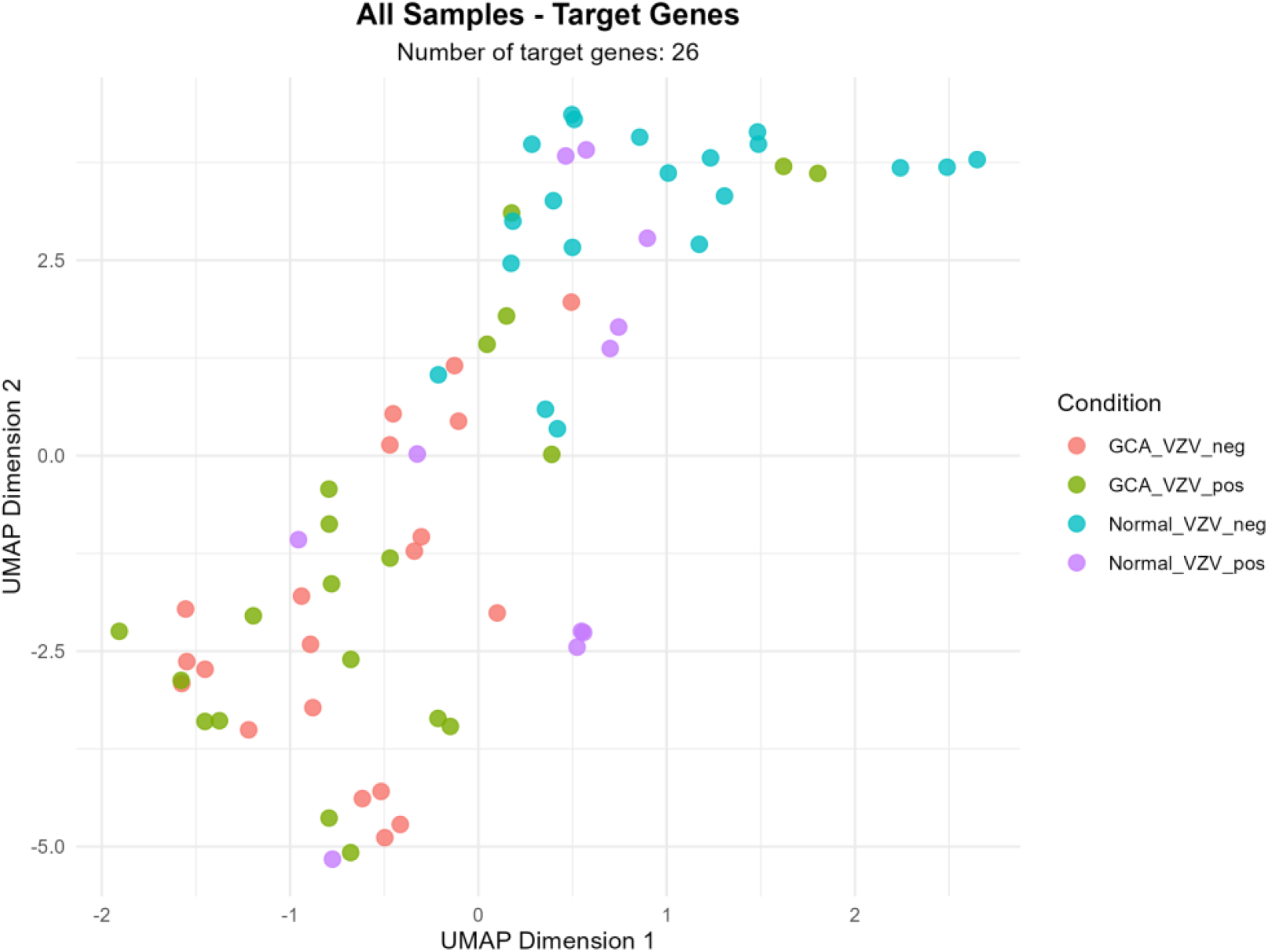
UMAP visualization of target genes in all samples (26 target genes in total). The horizontal axis is UMAP Dimension 1, and the vertical axis is UMAP Dimension 2; the sample grouping method is consistent with Figure 1, i.e., divided into four groups: GCA_VZV_neg, GCA_VZV_pos, Normal_VZV_neg and Normal_VZV_pos, and samples in each group are distinguished by differential markers.

**Table 12.**
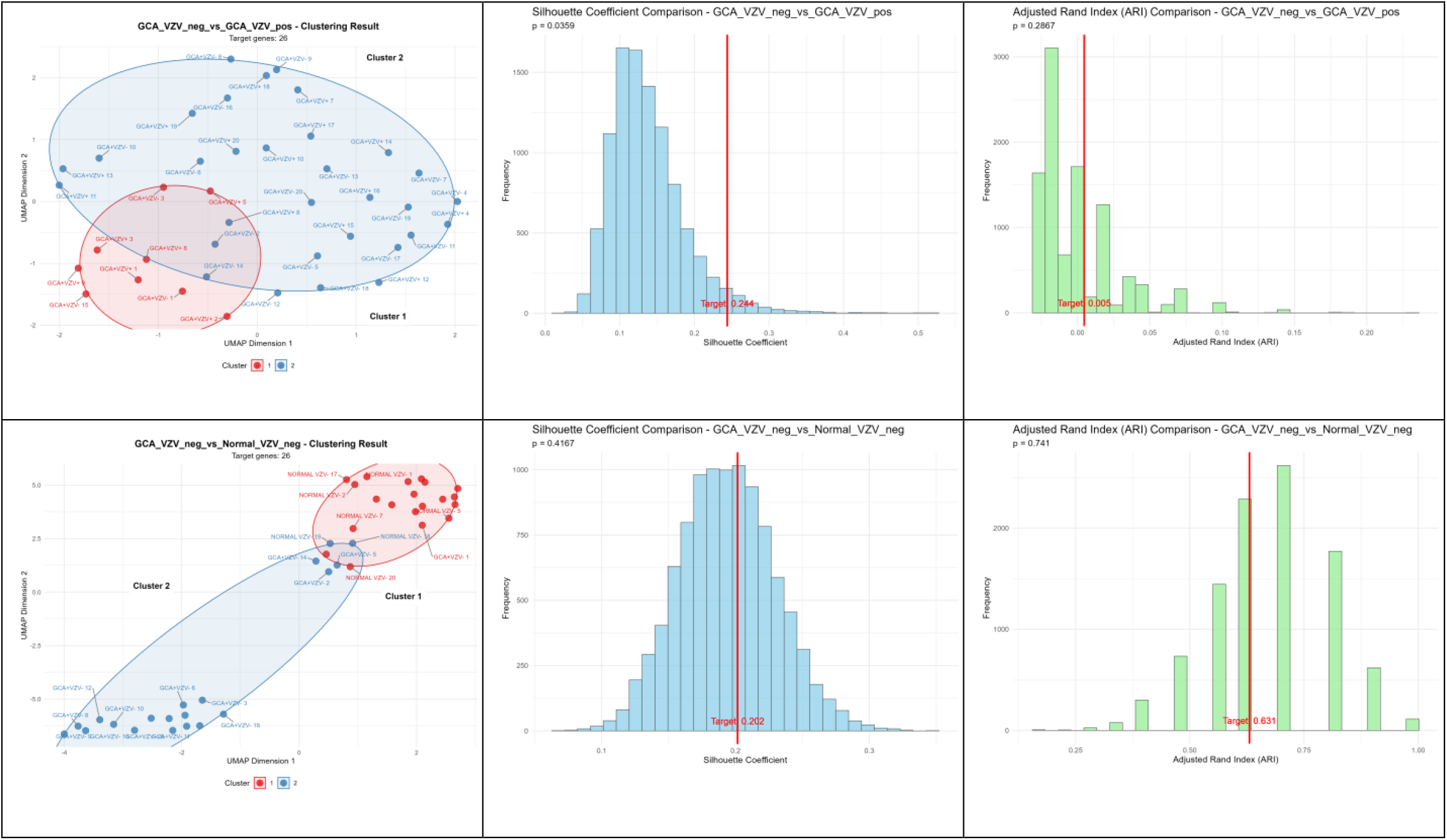
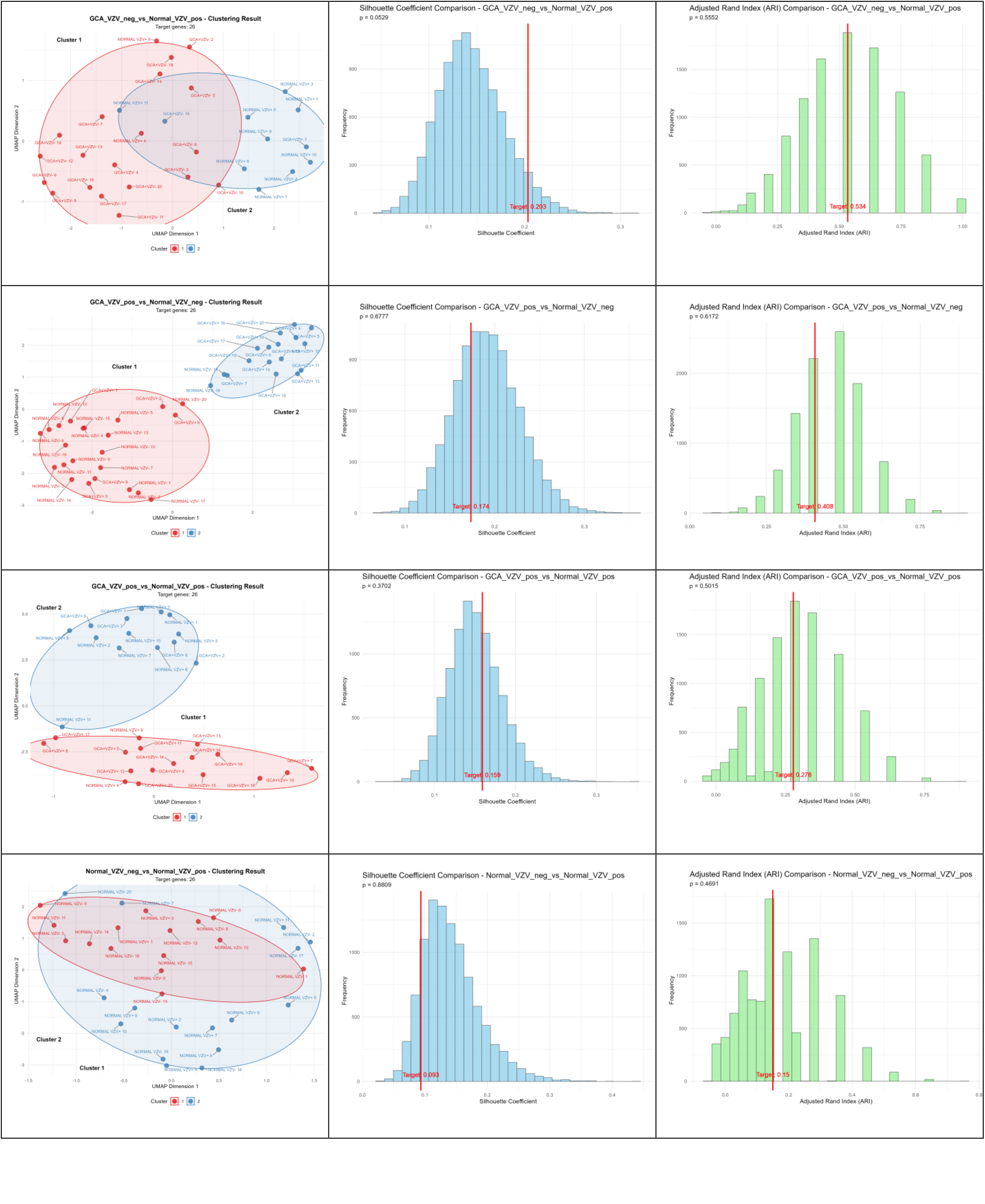
Summary table of clustering-related analysis subfigures based on VZV-labeled data. This table contains three columns of subfigures: the first column is UMAP clustering result subfigures, showing the UMAP clustering spatial distribution of samples under different experimental conditions; the second column is Silhouette Coefficient comparison subfigures, presenting the distribution characteristics of Silhouette Coefficients of sample clustering in different groups and the corresponding statistical test P-values; the third column is Adjusted Rand Index comparison subfigures, showing the distribution characteristics of Adjusted Rand Indexes of sample clustering in different groups and the corresponding statistical test P-values. Note: Due to table space limitations, subfigures related to affinity matrix and spectral clustering projection are omitted; since the data carries VZV labels, an Adjusted Rand Index analysis module is additionally added to more accurately evaluate the consistency of clustering results.

***Special Note: From the above analysis results, it can be observed that the Silhouette Coefficient is quite high, while the p-value of the Silhouette Coefficient comparison is not high; the Adjusted Rand Index is quite high, while the p-value of the Adjusted Rand Index comparison is not high. This implies that most genes in the transcriptome can distinguish between the two groups, which is obviously inconsistent with reality. Therefore, it is considered that there may be a situation where batch effects overlap with grouping. However, from the supervised clustering of GCA_VZV_neg VS GCA_VZV_pos, we can still see clues of the association between VZV and GCA. Additionally, we can also observe that the p-value of the Silhouette Coefficient comparison is correlated with the p-value of the Adjusted Rand Index comparison. In conclusion, in view of the batch effect issue of this dataset, it is still necessary to interpret the analysis results with caution.

### Discussion

Most of the studies cited above did not account for varicella-zoster virus (VZV) infection a priori; they are independent studies unrelated to VZV, can be regarded as small-sample cohort studies, and are effective in minimizing selection bias. The interacting proteins between VZV and humans can cluster GCA samples, and the clustering effect is stronger than that of most random gene sets. This indicates that the VZV-human interaction proteins are distinctive and that the probability of such clustering results being caused by outlier samples is extremely low. Additionally, the clustering effect of the control group is not significantly better than that of random gene sets, which indicates that the effect is not driven by the synergistic effect of proteins or genes, and rules out the possibility that the effect is dominated by highly expressed genes or abundant proteins. Therefore, this clustering result strongly suggests that VZV is associated with the pathogenesis of some giant cell arteritis (GCA) cases.

Two large-scale cohort studies suggest a relatively weak association between VZV and GCA. Rennie L Rhee et al ^[28]^ conducted a nested case-control study using UK electronic databases. They examined the association between prior infections (especially herpes zoster) and incident GCA in a population-based cohort. Rennie L Rhee et al found that antecedent infection was associated with GCA onset, though infection may be a secondary determinant of overall GCA risk. The study concluded that the moderate association between herpes zoster and GCA implies that herpes zoster infection is unlikely to play a major causal role in GCA pathogenesis. Bryant R England et al. ^[29]^performed a retrospective cohort study using two large, independent US administrative datasets: Medicare 5% and Truven Health Analytics MarketScan. Herpes zoster (HZ) events (both complicated and uncomplicated) and GCA were identified through physician visit records or hospital discharge records. Antiviral therapy and vaccination status were determined from prescription claims and medication codes. Bryant R England et al demonstrated an increased risk of GCA following HZ events. However, they noted that HZ events are less frequent in GCA patients, suggesting that HZ events may only have a meaningful impact on GCA risk in a small subset of patients. This finding aligns well with our study, which indicates that VZV is only involved in the pathogenesis of a subset of GCA cases. Furthermore, while such large-scale cohort studies are efficient and expansive, they are prone to limitations including missing clinical details, documentation bias, and incomplete follow-up—factors that may contribute to the observed weak association between VZV and GCA.

Could confounding factors exist that make certain patients both more susceptible to GCA and more likely to be infected with VZV? Some of the samples used in our clustering analysis were from patients with active GCA who had not received glucocorticoid (GC) treatment or had just initiated it, presenting with typical clinical symptoms and elevated inflammatory markers. Thus, the risk of VZV infection due to immunosuppression was low in these cases. A Mendelian randomization study ^[30]^ confirmed a causal association between genes related to primary immunodeficiency and GCA triggered by VZV reactivation. This association satisfies the exclusivity and independence assumptions, meaning that these genes induce vascular inflammation specifically by affecting VZV immune regulation pathways, rather than being driven by other confounding factors or reverse causality.

GCA is a complex disease, and ideal research models for it are lacking. The pathogenesis of VZV-induced GCA may be a long-term process. Animal models (e.g., mice and other model organisms) differ significantly from humans in terms of physiology, metabolism, lifespan, and immune system, making it difficult to fully simulate the genetic background and environmental influences of this complex human disease. Although in vitro models such as cell line cultures and organoids are useful, they cannot fully replicate the complex in vivo microenvironment (e.g., cell-cell interactions, immune regulation, and organ-system crosstalk). Ethical restrictions on human research prevent invasive experiments or environmental variable control in humans, so research primarily relies on observational studies (which are susceptible to confounding bias) or intervention studies with limited sample sizes. Therefore, randomized controlled trials (RCTs) of antiviral therapy in GCA patients may be the most useful approach to ultimately determine whether VZV contributes to the pathogenesis of some GCA cases. Our observations indicate that the interacting proteins between VZV and humans have the potential to distinguish VZV-related GCA cases, which provides a basis for screening GCA patients who may benefit from antiviral therapy.

Our study also has limitations. First, the sample size is small. Additionally, while unsupervised clustering analysis is a commonly used exploratory tool for identifying potential patterns, subgroups, or structures in data (e.g., distinguishing different patient subtypes, cell subsets, or gene expression patterns), it has significant limitations. In unsupervised learning, there are no pre-defined true labels to evaluate the quality of clustering results. Unsupervised clustering is a tool for generating hypotheses rather than verifying them; its results require cautious interpretation and should be treated as hypotheses to be validated.

### Future Outlook

In the future, first, new studies with larger sample sizes are needed to validate the above findings. Second, it will be necessary to include VZV infection-related tests in all future GCA studies. It is also important to continue exploring VZV infection-related biological markers in subsets of GCA patients. Antiviral therapy RCTs may serve as the “final verdict” to confirm whether VZV is an etiological factor for some GCA cases. Emerging technologies such as spatial single-cell technology may help us explore the molecular network associations between VZV and GCA. As Ellen F Foxman ^[31]^ stated: “If we understand the mechanisms by which host-virus interactions cause complex diseases, we may be able to develop new approaches to address these diseases. The prevalence of autoimmune and inflammatory diseases is on the rise; understanding the role of the virome may help clarify the causes of this epidemiological trend and identify intervention strategies. Furthermore, if we can identify altered host-virus interactions that promote the pathogenesis of complex diseases, it may be possible to design therapies that shift these interactions toward the responses seen in non-susceptible individuals, thereby protecting susceptible populations. Future research on host-virome interactions is expected to provide new methods for the diagnosis and treatment of complex diseases. “

## REFERENCES

[1] Helmick CG, Felson DT, Lawrence RC, et al. Estimates of the prevalence of arthritis and other rheumatic conditions in the United States. Part I. Arthritis Rheum. 2008;58(1):15–25. doi:10.1002/art.23177

[2] Gonzalez-Gay MA, Vazquez-Rodriguez TR, Lopez-Diaz MJ, et al. Epidemiology of giant cell arteritis and polymyalgia rheumatica. Arthritis Rheum. 2009;61(10):1454–1461. doi:10.1002/art.24459

[3] Salvarani C, Pipitone N, Versari A, Hunder GG. Clinical features of polymyalgia rheumatica and giant cell arteritis. Nat Rev Rheumatol. 2012;8(9):509–521. doi:10.1038/nrrheum.2012.97

[4] Tomasson G, Peloquin C, Mohammad A, et al. Risk for cardiovascular disease early and late after a diagnosis of giant-cell arteritis: a cohort study. Ann Intern Med. 2014;160(2):73–80. doi:10.7326/M12-3046

[5] Aviña-Zubieta JA, Bhole VM, Amiri N, Sayre EC, Choi HK. The risk of deep venous thrombosis and pulmonary embolism in giant cell arteritis: a general population-based study. Ann Rheum Dis. 2016;75(1):148–154. doi:10.1136/annrheumdis-2014-205665

[6] Weyand CM, Goronzy JJ. Immune mechanisms in medium and large-vessel vasculitis. Nat Rev Rheumatol. 2013;9(12):731–740. doi:10.1038/nrrheum.2013.161

[7] Agger WA, Deviley JA, Borgert AJ, Rasmussen CM. Increased Incidence of Giant Cell Arteritis After Introduction of a Live Varicella Zoster Virus Vaccine. Open Forum Infect Dis. 2020;8(2):ofaa647. Published 2020 Dec 30. doi:10.1093/ofid/ofaa647

[8] Elkind MS. The varicella zoster virus vasculopathies: clinical, CSF, imaging, and virologic features. Neurology. 2009;72(11):1028–130. doi:10.1212/01.wnl.0000339389.10848.4a

[9] Mitchell BM, Font RL. Detection of varicella zoster virus DNA in some patients with giant cell arteritis. Invest Ophthalmol Vis Sci. 2001;42(11):2572–2577.

[10] Nagel MA, White T, Khmeleva N, et al. Analysis of Varicella-Zoster Virus in Temporal Arteries Biopsy Positive and Negative for Giant Cell Arteritis. JAMA Neurol. 2015;72(11):1281–1287. doi:10.1001/jamaneurol.2015.2101

[11] Gilden D, White T, Khmeleva N, Katz BJ, Nagel MA. Blinded search for varicella zoster virus in giant cell arteritis (GCA)-positive and GCA-negative temporal arteries. J Neurol Sci. 2016;364:141–143. doi:10.1016/j.jns.2016.03.020

[12] Gilden D, White T, Khmeleva N, et al. Prevalence and distribution of VZV in temporal arteries of patients with giant cell arteritis. Neurology. 2015;84(19):1948–1955. doi:10.1212/WNL.0000000000001409

[13] Gilden D, White T, Boyer PJ, et al. Varicella Zoster Virus Infection in Granulomatous Arteritis of the Aorta. J Infect Dis. 2016;213(12):1866–1871. doi:10.1093/infdis/jiw101

[14] Kennedy PG, Grinfeld E, Esiri MM. Absence of detection of varicella-zoster virus DNA in temporal artery biopsies obtained from patients with giant cell arteritis. J Neurol Sci. 2003;215(1-2):27–29. doi:10.1016/s0022-510x(03)00167-9

[15] Nordborg C, Nordborg E, Petursdottir V, et al. Search for varicella zoster virus in giant cell arteritis. Ann Neurol. 1998;44(3):413–414. doi:10.1002/ana.410440323

[16] Alvarez-Lafuente R, Fernández-Gutiérrez B, Jover JA, et al. Human parvovirus B19, varicella zoster virus, and human herpes virus 6 in temporal artery biopsy specimens of patients with giant cell arteritis: analysis with quantitative real time polymerase chain reaction. Ann Rheum Dis. 2005;64(5):780–782. doi:10.1136/ard.2004.025320

[17] Helweg-Larsen J, Tarp B, Obel N, Baslund B. No evidence of parvovirus B19, Chlamydia pneumoniae or human herpes virus infection in temporal artery biopsies in patients with giant cell arteritis. Rheumatology (Oxford). 2002;41(4):445–449. doi:10.1093/rheumatology/41.4.445

[18] Rodriguez-Pla A, Bosch-Gil JA, Echevarria-Mayo JE, et al. No detection of parvovirus B19 or herpesvirus DNA in giant cell arteritis. J Clin Virol. 2004;31(1):11–15. doi:10.1016/j.jcv.2004.05.003

[19] Verdijk RM, Ouwendijk WJD, Kuijpers RWAM, Verjans GMGM. No Evidence of Varicella-Zoster Virus Infection in Temporal Artery Biopsies of Anterior Ischemic Optic Neuropathy Patients With and Without Giant Cell Arteritis. J Infect Dis. 2021;223(1):109–112. doi:10.1093/infdis/jiaa566

[20] Buckingham EM, Foley MA, Grose C, et al. Identification of Herpes Zoster-Associated Temporal Arteritis Among Cases of Giant Cell Arteritis. Am J Ophthalmol. 2018;187:51–60. doi:10.1016/j.ajo.2017.12.017

[21] Solomon IH, Docken WP, Padera RF Jr. Investigating the Association of Giant Cell Arteritis with Varicella Zoster Virus in Temporal Artery Biopsies or Ascending Aortic Resections. J Rheumatol. 2019;46(12):1614–1618. doi:10.3899/jrheum.180912

[22] Sammel AM, Smith S, Nguyen K, et al. Assessment for varicella zoster virus in patients newly suspected of having giant cell arteritis. Rheumatology (Oxford). 2020;59(8):1992–1996. doi:10.1093/rheumatology/kez556

[23] Nitzan I, Shemesh N, Kubovsky S, Shalmov T, Levy J, Amer R. Incidence of Giant Cell Arteritis Following Herpes Zoster Ophthalmicus: A Multicenter Retrospective Cohort Study. Am J Ophthalmol. Published online May 23, 2025. doi:10.1016/j.ajo.2025.05.019

[24] Cunningham KY, Hur B, Gupta VK, et al. Plasma proteome profiling in giant cell arteritis. Ann Rheum Dis. 2024;83(12):1762–1772. Published 2024 Nov 14. doi:10.1136/ard-2024-225868

[25] Estupiñán-Moreno E, Ortiz-Fernández L, Li T, et al. Methylome and transcriptome profiling of giant cell arteritis monocytes reveals novel pathways involved in disease pathogenesis and molecular response to glucocorticoids. Ann Rheum Dis. 2022;81(9):1290–1300. Published 2022 Aug 11. doi:10.1136/annrheumdis-2022-222156

[26] Hur B, Koster MJ, Jang JS, Weyand CM, Warrington KJ, Sung J. Global Transcriptomic Profiling Identifies Differential Gene Expression Signatures Between Inflammatory and Noninflammatory Aortic Aneurysms. Arthritis Rheumatol. 2022;74(8):1376–1386. doi:10.1002/art.42138

[27] Bubak AN, Mescher T, Mariani M, et al. Targeted RNA Sequencing of Formalin-Fixed, Paraffin-Embedded Temporal Arteries From Giant Cell Arteritis Cases Reveals Viral Signatures. Neurol Neuroimmunol Neuroinflamm. 2021;8(6):e0178. Published 2021 Sep 7. doi:10.1212/NXI.0000000000001078

[28] Rhee RL, Grayson PC, Merkel PA, Tomasson G. Infections and the risk of incident giant cell arteritis: a population-based, case-control study. Ann Rheum Dis. 2017;76(6):1031–1035. doi:10.1136/annrheumdis-2016-210152

[29] England BR, Mikuls TR, Xie F, Yang S, Chen L, Curtis JR. Herpes Zoster as a Risk Factor for Incident Giant Cell Arteritis. Arthritis Rheumatol. 2017;69(12):2351–2358. doi:10.1002/art.40236

[30] Wang H, Chen G, Gong Q, Wu J, Chen P. Primary immunodeficiency-related genes and varicella-zoster virus reactivation syndrome: a Mendelian randomization study. Front Immunol. 2024;15:1403429. Published 2024 Aug 26. doi:10.3389/fimmu.2024.1403429

[31] Foxman EF, Iwasaki A. Genome-virome interactions: examining the role of common viral infections in complex disease. Nat Rev Microbiol. 2011;9(4):254–264. doi:10.1038/nrmicro2541

